# Crystal structure of the Rubella virus protease reveals a unique papain-like protease fold

**DOI:** 10.1101/2022.04.15.488536

**Authors:** Ezekiel Ze Ken Cheong, Jun Ping Quek, Liu Xin, Chaoqiang Li, Jing Yi Chan, Chong Wai Liew, Yuguang Mu, Jie Zheng, Dahai Luo

**Author notes:** These authors contributed equally to this work (Ezekiel Ze Ken Cheong, Jun Ping Quek).

## Abstract

Rubella is well-controlled due to an effective vaccine, but outbreaks are still occurring without any available antiviral treatments. There is still much to learn about the rubella virus (RUBV) papain-like protease (RubPro) that could be a potential drug target. This protease is crucial to RUBV replication, cleaving the non-structural polyprotein p200 into 2 multi-functional proteins, p150 and p90. Here we report a novel crystal structure of RubPro at 1.64 Å resolution. It has a similar catalytic core structure to that of SARS-CoV-2 and foot-mouth-disease virus (FMDV) proteases. RubPro has well-conserved sequence motifs that are also found in its newly discovered *Rubivirus* relatives. The RubPro construct was shown to have protease activity in *trans* against a construct of RUBV protease-helicase and fluorogenic peptide. A protease-helicase construct was also cleaved in *E. coli* expression. RubPro was demonstrated to possess deubiquitylation activity, suggesting a potential role of RubPro in modulating the host’s innate immune responses. The structural and functional insights of the RubPro will advance our current understanding of its function and point to more structure-based research into the RUBV replication machinery, in hopes of developing antiviral therapeutics in the future.

## Introduction

Rubella is an infectious disease that is well-characterised by rashes [1]. It is often confused with measles that also cause rashes. Confoundingly, rubella is also known as German measles, and measles is also known as rubeola [2]. However, rubella and measles are different diseases caused by different viruses. The rubella virus (RUBV) is the aetiological agent of the Rubella disease and belongs to the genus, *Rubivirus*, of the newly created family, *Matonaviridae* [3]. Rubella infection during the first trimester of pregnancy can result in miscarriage or congenital rubella syndrome (CRS). CRS is characterised by foetal cataracts, deafness, heart defects and global developmental delay [4]. Infection at the early stages of pregnancy typically has the worst prognosis [5]. There is currently no treatment for CRS apart from symptomatic treatment [6]. Before the rubella vaccine was developed in 1969, rubella epidemics occurred every 6-9 years [7]. Modern rubella vaccines utilise the RA27/3 strain [8], a strain of RUBV obtained from an aborted foetus infected with the virus [9]. The vaccine is typically administered in a combination of measles, mumps and rubella (MMR) vaccines. With 97% effectiveness [10], the vaccine has virtually eliminated rubella in more than 130 countries [11]. As such, there appears to be very little impetus to research deeply into its virology.

RUBV is a Group VI virus with a single-stranded, positive-sense RNA genome enclosed by an icosahedral capsid [4, 12]. The viral genome is around 10 kb in size and has the highest GC content of RNA viruses, at 70% [13]. The genome has a 5’ cap structure with a poly(A) tail at its 3’end. The 5’-proximal open reading frame (ORF) encodes the non-structural polypeptide p200, while the 3’-proximal ORF encodes the structural proteins, capsid and surface glycoproteins E1 and E2 (Fig 1A) [14]. The non-structural polyprotein p200 is then processed into two non-structural proteins, p150 and p90 [15]. The p150 protein consists of a methyltransferase (MTase) and protease domain [16], while the p90 protein has both helicase and RNA-dependent RNA polymerase (RdRp) domain [17]. These non-structural proteins are crucial to RNA viruses for replication [18] and polyprotein processing [19].

**Figure 1.**
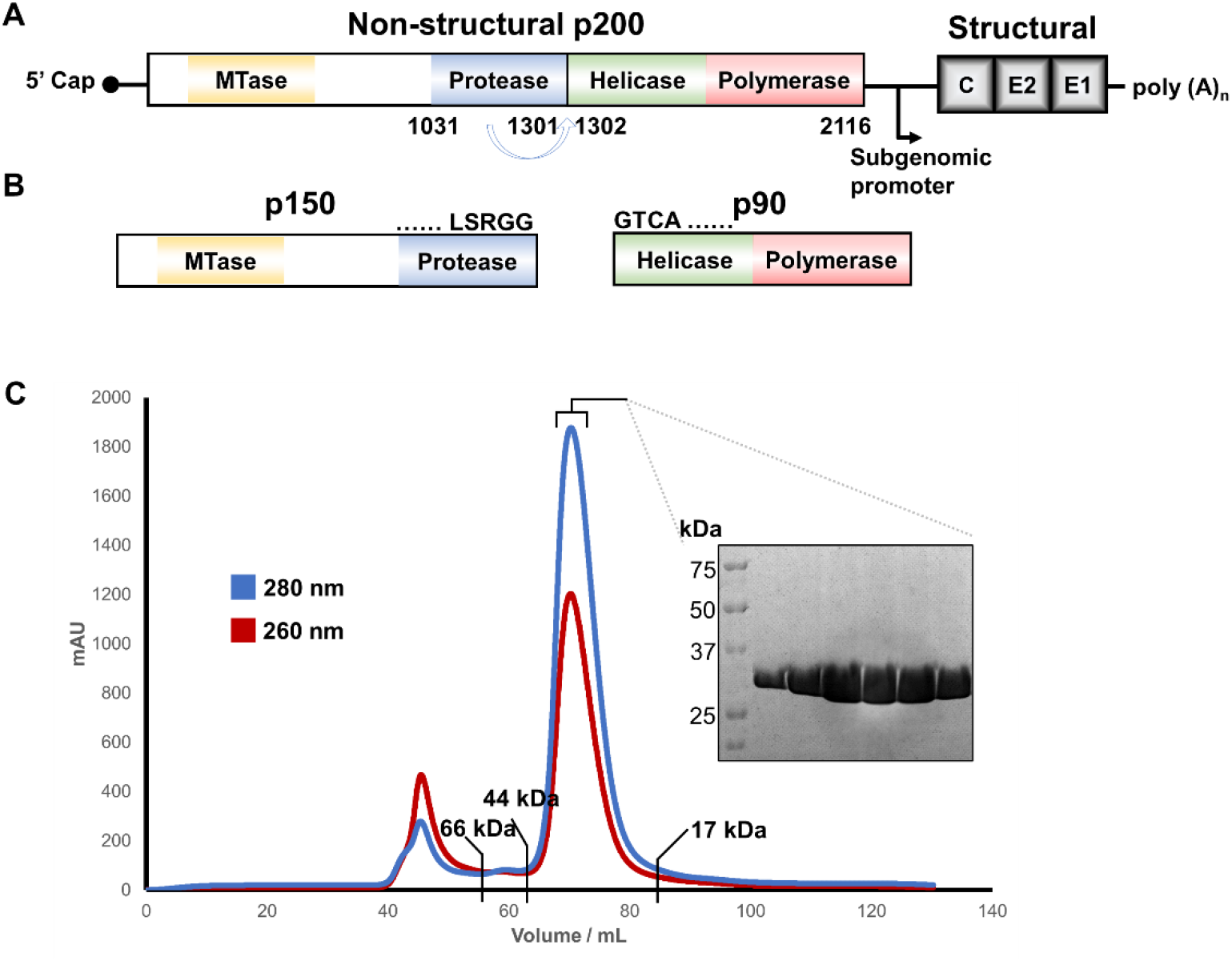
RUBV genome and RUBV protease function and expression. (**A**) Schematic representation of the RUBV genome. The 5’ ORF encodes for the non-structural protein p200 while the 3’ ORF encodes the structural proteins, capsid (C) and surface glycoproteins E1 & E2. The enzymatic domains of the non-structural polypeptide are labelled, with the protease domain highlighted in blue. (**B**) The protease domain located within p150 will cleave the non-structural p200 into p150 and p90. The cleavage site is at SRGG/GTCA. (**C**) The gel filtration profile of RubPro on Superdex 75 16/60 column. column (GE Healthcare). SDS-PAGE analysis of the RubPro with an expected molecular weight of 30.3 kDa. The elution volumes of 3 other standard proteins are labelled as reference

The RUBV protease, RubPro, while in the p200, cleaves the polyprotein between residues G1301 and G1302 [20]. This cleavage, at SRGG/GTCA, is found at the C-terminal of RubPro in p150 and the N-terminal of the helicase domain of p90 (Fig 1B) [21]. Based on computer alignments, RubPro is predicted to be a papain-like cysteine protease (PCP). PCPs form a large family among cysteine proteases and can be found in plants, viruses and parasites [22]. Coronaviruses and alphaviruses utilise PCPs for polypeptide processing and immune evasion mechanisms [23, 24]. Severe acute respiratory syndrome coronavirus 2 (SARS-CoV-2) papain-like protease (PLpro) [25] and foot-and-mouth disease (FMDV) PCP [26, 27] cleaves inflammatory ubiquitin and other ubiquitin-like antiviral proteins, such as interferon-stimulated gene 15 (ISG15), to modulate host innate immunity responses [28]. The functional importance of PCPs for viral replication and survival makes them viable drug targets for antiviral treatment [29]. RubPro has residues C1152 and H1273 as the catalytic dyad [30], confirmed via site-directed mutagenesis [31]. RubPro requires divalent ions like Zn^2+^, Co^2+^ and Cd^2+^ for protease activity [32]. Additionally, the Ca^2+^-dependent association of calmodulin with RubPro is necessary for the proteolytic activity of RubPro [33]. Using FMDV PCP as a reference, homology modelling has been used to partially model the RubPro structure [34]. RubPro contains a Ca^2+^-binding EF-hand domain which plays a role in maintaining the structure of RubPro [34]. The cysteine residues are arranged close to Zn^2+^, and this Zn^2+^-coordination also contributes to structural stability [35]. The structure of RubPro has not been solved, and its enzymatic activity or inhibition has not been characterised quantitatively.

Scientific knowledge of RUBV is relatively superficial compared to other viruses, likely due to the effective vaccine reducing the need for deeper research. The only proteins of RUBV with solved structures are the structural proteins E1 [36] and capsid protein [37]. The enzymatic and essential non-structural proteins’ structures and interactions are not well-understood or adequately characterised. While only RUBV is found naturally in humans [38], two newly discovered *Rubivirus*, Ruhugu virus (RUHV) and Rustrela virus (RUSV) have crossed barriers across other mammalian host species. This raised concerns about the potential zoonotic transmission of RUBV-like viruses into human hosts [39]. Furthermore, despite an effective RUBV vaccine, there are still rubella outbreaks occurring due to gaps in national immunisation programmes. Japan experienced three rubella outbreaks during the last decades [40, 41]. There were outbreaks in Poland [42] and Romania [43] in 2012 as well. Once an outbreak manages to break the first line of defence, which is vaccine coverage, there are no antiviral treatments available to combat the virus and the disease. Effective antiviral therapeutics requires a fundamental understanding of viral replication processes. Many successful drugs inhibit proteins of the replication machinery, such as proteases [44, 45] and RNA polymerases [46-48]. The lack of knowledge about RUBV highlights a need for deeper research into its replication to potentially develop antiviral treatments, as well as to expand the knowledge of virology that could enlighten the replication and pathogenesis mechanisms of other viruses.

In this study, the crystal structure of RubPro to a resolution of 1.64 Å is reported. This novel structure provides the molecular basis for the substrate recognition and characterisation of the active site for potential inhibitor studies. Next, sequence alignment was conducted with RUHV and RUSV proteases. RubPro was also structurally aligned with other viral PCPs deposited in the Protein Data Bank (PDB) for further structural analysis. The protease’s activity was also characterised using RUBV protease-helicase as the natural substrate as well as using fluorogenic peptides. RubPro was shown to have K48-linkage-specific deubiquitylation activity. This study provides structural and functional insight into RubPro to facilitate antiviral development against RUBV, as well as to expand the field of knowledge of viral proteins.

## Materials and methods

### Designs of protein constructs

Cloning and expression tests of various proteins were carried out by the Protein Production Platform (PPP) at Nanyang Technological University (NTU).

RubPro, RubPro C1152A mutant and RUBV protease-helicase (RubProHel) C1152A mutant were cloned in pSUMO-LIC vector, which encodes an N-terminal His-Sumo tag that is cleavable by Sumo protease. RubPro consisted of RUBV residues 1021-1301 and RubProHel consisted of RUBV residues 1021-1616. C1152A mutants were generated by site-directed mutagenesis of the respective wild type (WT) constructs.

### Mutagenesis of RubPro and RubProHel

C1152A mutagenesis were performed, for both RubPro and RubProHel using Q5 Site-directed Mutagenesis Kit (New England BioLabs). The forward and reverse primer used were TCCCAACACTGCGTGGCTGAGAGCCGCCG and TCGAGTTCCCCGCCCCTT respectively. Mutagenesis for both constructs were conducted as per the manufacturer’s protocol.

### Expression and purification of protein constructs

The protein expression plasmids were transformed into *Escherichia coli* Rosetta 2 DE3 strain. The bacteria were cultured at 37°C in LB broth supplemented with kanamycin and chloramphenicol. They were induced with 1 mM IPTG and cultured overnight at 18°C. Bacteria cells were harvested by spinning in a Sorvall LYNC 4000 centrifuge (Thermo Fisher Scientific) at 6,000 rpm for 15 min at 4°C.

The pellet was resuspended in lysis buffer containing 1x PBS, supplemented with 160 mM NaCl, 5 mM β-mercaptoethanol and 10% v/v glycerol. The sample was sonicated at 70% amplitude for 5 min at a 5s on / 5s off interval using a Vibra-Cell sonicator (Sonics). The cells were further lysed using PandaPLUS 2000 homogenizer (GEA). The sample was centrifuged in a Sorvall LYNC 4000 centrifuge at 20,000 rpm for 1 h at 4°C. The supernatant was collected and filtered with a 0.45 μm filter. The supernatant was subjected to 1.5 h of incubation with HisPur Ni-NTA Resin (Thermo Fisher Scientific) at 4°C for immobilised-metal affinity chromatography (IMAC). Next, the resin was washed with lysis buffer supplemented with 20 mM imidazole and eluted with lysis buffer with 300 mM imidazole. Fractions were analysed with SDS-PAGE. Chosen fractions were incubated with Sumo protease at 4°C, in dialysis with lysis buffer as the dialysate. Cleaved proteins were collected in reverse IMAC using Ni-NTA resin, where the flowthrough was collected and concentrated in concentrators (Merck Millipore) with appropriate molecular weight cut-off (MWCO) filter sizes. Samples then underwent size exclusion chromatography in an ÄKTA Pure fast protein liquid chromatography (FPLC) using HiLoad 16/600 Superdex 200 or Superdex 75 column (GE Healthcare Life Sciences) in GF-300 buffer (25 mM HEPES buffer at pH 7.5, 300 mM NaCl, 5% w/v glycerol and 2 mM DTT). Fractions were analysed with SDS-PAGE to determine purity. Pure fractions were then concentrated in concentrators of appropriate MWCO filter sizes. The purified proteins were then aliquoted, flash-frozen with liquid N_2_ and stored at - 80°C.

### Protein crystallisation

1 μL of RubPro at a final concentration of 16 mg/mL in a GF-300 buffer supplement with 2 mM TCEP, was mixed with 1 μL of reservoir solution containing 0.1 M HEPES pH 7.5 and 15% w/v polyethylene glycol 3,350, the mixture was subjected to crystallisation via hanging drop vapour diffusion method at 20°C. Crystals appeared after two days of incubation. The crystals were mounted onto a cryo-loop and cryoprotected with reservoir solution supplemented with 20% v/v glycerol, and flash-frozen in liquid N_2_.

### Data collection, structure determination and model building

Diffraction intensities of the native crystals were recorded on Dectris EIGER 16M detector at MX2 beamline at wavelength 1.284 Å at the Australian Synchrotron, Melbourne, Australia. Data processing was carried out using the XDS data processing package.

The structure of the RubPro was determined using the single-wavelength anomalous diffraction (SAD) method. Measurements of anomalous signals from Zn^2+^ were used to derive the positions of the heavy atom substructure. Crank2 [49] and BUCCANEER [50] software were used to calculate the initial phases and for automatic model building respectively. The structure was subjected to iterative rounds of refinement using the Phenix_refine program [51-53] and manual rebuilding using Coot [54]. Figures were generated using PyMOL (Schrödinger).

Data collection and refinement statistics can be found in Table S1.

### Sequence alignment with RUHV and RUSV proteases

The amino acid residue sequence of our RubPro construct was aligned with the sequences of the non-structural p200 polypeptides of RUHV and RUSV. The sequences were obtained from UniProtKB with the identifiers A0A7L5KV54 for RUHV and A0A7L5KV68 for RUSV. Alignments were done using Clustal Omega [55].

### Testing of RubProHel C1152A cleavage by RubPro

0.5 mg/mL of RubProHel C1152A was incubated with doubling concentrations of RubPro, from 0.125 mg/mL to 2 mg/mL, at room temperature for 18 h in cleavage buffer (25 mM HEPES buffer at pH 7.5, 150 mM NaCl, 5% w/v glycerol and 2 mM DTT). Samples were analysed analyzed by electrophoresis on a 15% SDS-PAGE gel and visualized by Coomassie blue staining.

### Enzyme concentration optimisation for protease assay

The assay was performed in a 96-well half-area black clear-bottom microplate (Corning). The reaction consists of varying concentrations of RubPro from 0.9 to 15 μM in assay buffer containing 25 mM Hepes pH 6.8 and 150 mM NaCl. The measurements were started by adding the Benzyloxycarbonyl-Arg-Leu-Arg-Gly-Gly-4-methylcoumaryl-7-amide (Z-RLRGG-MCA) peptide substrate (Peptide Institute Inc, Japan) at 50 μM. The relative fluorescence readings were measured using Synergy H1 Microplate Reader (BioTek) at 3-minute intervals over 1 hour with the excitation wavelength (λ_ex_) at 380 nm and emission wavelength (λ_em_) at 460 nm. The assays were conducted as triplicates at 37 °C.

### Buffer pH optimisation for protease assay

The assay was performed in a 96-well half-area black clear-bottom microplate (Corning). The reaction consists of 5 μM of RubPro in assay buffer containing 50 mM potassium phosphate buffer with varying pH from 5.8 to 7.8. The measurements were started by adding the Z-RLRGG-MCA peptide substrate (Peptide Institute Inc, Japan) at 50 μM. The relative fluorescence readings were measured using Synergy H1 Microplate Reader (BioTek) at 3-minute intervals over 1 hour with the excitation wavelength (λ_ex_) at 380 nm and emission wavelength (λ_em_) at 460 nm. The assays were conducted as triplicates at 37 °C.

### Protease assay

Z-RLRGG-MCA substrate with starting concentration of 2.5 mM was serially diluted two times in assay buffer (50 mM potassium phosphate buffer pH 6.6) and added to Corning 96 Well black plates containing 5 μM RubPro or RubPro C1152A mutant diluted in the same buffer. The relative fluorescence readings were measured using Synergy H1 Microplate Reader (BioTek) at 3-minute intervals over 1 hour with the excitation wavelength (λ_ex_) at 380 nm and emission wavelength (λ_em_) at 460 nm. The assays were carried out as triplicates at 37 °C. To determine the amount of AMC released, a standard AMC curve was plotted with various concentrations of AMC. Initial velocities were calculated using the linear regression function in the GraphPad Prism software. Data were analysed and plotted using the Michaelis-Menten equation with GraphPad Prism 8 for Windows (GraphPad Software, San Diego California USA).

The assay was repeated using another chemically synthesised peptide substrate, Acryl-LSRGG-AMC (GenScript, Hong Kong).

### Structural prediction of RubPro using Alphafold2 web server

Structural prediction of RubPro was performed using the Alphafold2 Colab web server [57].

### Docking and MD simulations

Isolated peptides, Ace-Leu-Ser-Arg-Gly-Gly-Gly-Nme (LSRGGG) and Ace-Arg-Leu-Arg-Gly-Gly-Gly-Nme(RLRGGG), were simulated in a water box with size of 3.82 nm×3.82 nm×3.82 nm for 500 ns. From the second 250 ns trajectories, one structure of every 100 ps was chosen (in total 2500 structures for each peptide) to dock onto the protein structure near the active side (C1152 and H1273). The quick autodock vina tool [56] was used for docking by freezing all the peptide conformation. These docked structures were ranked based on the distance between the amide nitrogen atom of the third glycine and the center of C1152 and H1273, the catalytic dyad. Five poses of each peptide were chosen with the distance (above mentioned) less than 0.5 nm. Molecular dynamics simulations were performed in the explicit water box for the chosen protein-peptide complexes. In most of the simulations, the peptide moves away from the active site. Only two systems of LSRGGG show stable binding of the peptide. The last 2 glycine residues were found forming hydrogen-bonds with the residue near the catalytic dyad, however, the distance between the amide nitrogen atom of the third glycine and the center of C1152 and H1273 is still far, around 0.6 nm. Here we made a structure manipulation: based on the binding pose found in the simulation, we shifted the peptide backbone towards the active side by 2 residues by superimposing the first glycine (G4) to the last glycine (G6). The new complex structure was used to perform MD simulation in the explicit water box to equilibrate. In this way, the protein-LSRGGG complex structure was obtained. The initial protein-RLRGGG structure was got based on the protein-LSRGGG structure and mutating LSRGGG to RLRGGG.

Parameters of the protein and peptides were based on the AMBER99SB-ILDN force field [57] and the system was solvated with TIP3P [58] water molecules and counterions were added to neutralize the system. The molecular dynamics (MD) simulations were performed using GROMACS [59] 5.1.2 software. The LINCS [60] algorithm was used to constrain bonds between heavy atoms and hydrogen to enable a timestep of 2 fs. A 1.2nm cutoff was used for van der waals interaction and short-range electrostatic interactions calculations, and the Particle Mesh Ewald method was implemented for long range electrostatic calculations. Simulation temperature was maintained at 300K using a V-rescale thermostat [61] and 1bar pressure using Parrinello-Rahman [62] barostat.

### Binding energy calculations

To analyse the behaviour of interactions between the protein and the peptides, we calculated the binding energies between them using the MM-GBSA (Molecular Mechanics Generalized Born Surface Area) method[63]. The entropy was not calculated:

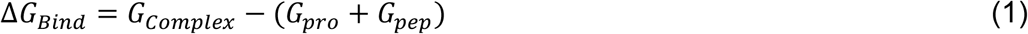

Where, G_Complex_, G_pro_ and G_pep_ is the free energy of complex, the protein, and the peptide, respectively. Free energy (ΔG) of each state was calculated as follows:

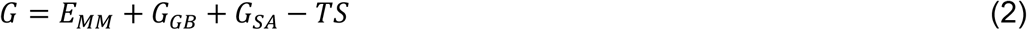

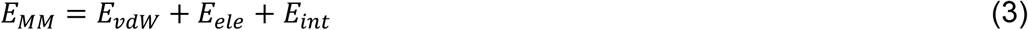

Where E_MM_ is the molecular mechanical energy, G_GB_ is the polar contribution towards solvation energy calculated by Generalized Born (GB) method respectively. G_SA_ is the contribution from the nonpolar terms towards solvation energy, and TS is the entropic contribution of the system. E_MM_ was obtained by summing contributions from the electrostatic energy (E_ele_), the van der Waal energy (E_vdw_), and the internal energy including bond, angle, and torsional angle energy terms (E_int_) using the same force field as that of MD simulations. G_GB_ was calculated with Onufriev’s method [64]. G_SA_ in equation 2 is proportional to the solvent accessible surface area (SASA) and was computed by molsurf module:

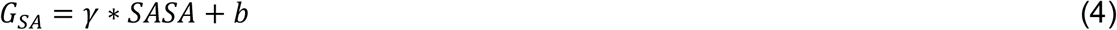

Where, the surface tension proportionality constant (γ) was set to 0.0072 kcal/mol/Å^2^, while the free energy of nonpolar solvation for a point solute (b) was set to a default value.

### In vitro assay for deubiquitylating and deISGylating

The protein substrates ubiquitination was prepared as previously described [65]. Purified RubPro wild-type protein or mutant (C1152A) protein was incubated with 8 μM of the substrate (K63-triubiquitin, K48-triubiquitin and M1-triubiquitin(linear) chains and ISG15-HIS) at 25°C for 5 to 30 min in 10 μl of reaction buffer containing 50 mM Tris-HCl, pH 7.5, 150 mM NaCl, 5% glycerol. The reaction was quenched by the addition of SDS sample buffer, analyzed by electrophoresis on a 15% SDS-PAGE gel and visualized by Coomassie blue staining.

## Results

### Structure determination of RubPro

The novel crystal structure of RubPro was determined at a resolution of 1.64 Å collected at Zn K absorption edge of 9.7 KeV, using the RubPro domain of the P150 (Residues 1021-1301) (Fig 2 and Fig S1). The RubPro adopts a unique right-hand architecture with two Zn^2+^ binding sites and a catalytic Cys-His dyad. The fingers, palm and thumb domains correspond to the N-terminal, middle region and the C-terminal of RubPro respectively. The RubPro fingers domain consists of 3 short parallel β-sheets and 4 α-helices. The C1034, C1037, C1040 and H1088 coordinate a Zn^2+^ ion near β3. Next, the palm domain consists of 4 α-helices, with the catalytic C1152 found on α5. The residues C1175, C1178, C1225 and C1227 coordinate a Zn^2+^ ion near α6. Lastly, the thumb domain consists of 5 β-sheets with the other catalytic residue, H1273, on β7.

**Figure 2.**
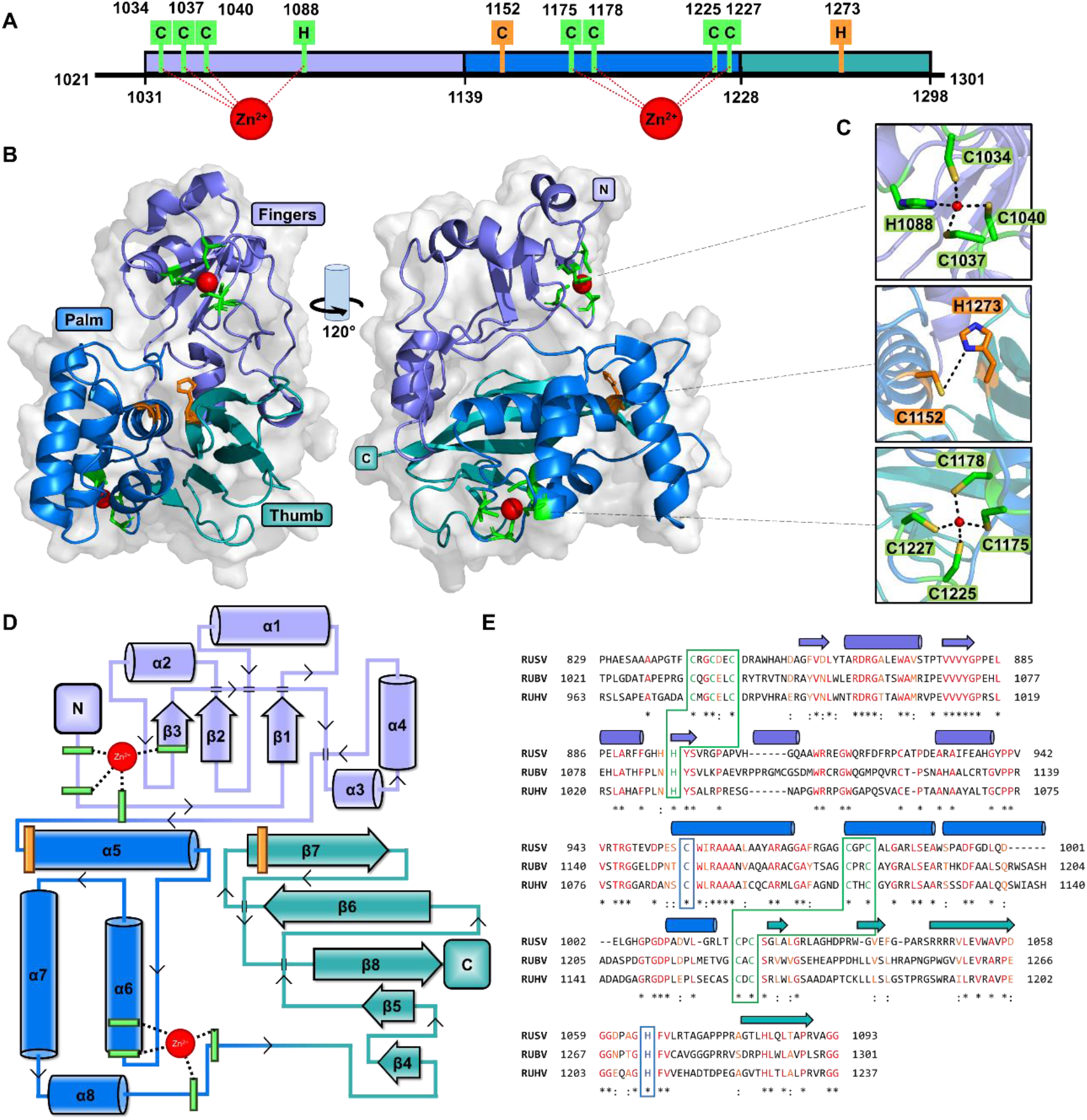
Crystal structure of RubPro. (**A**) Schematic representation of RubPro with residues of interest being highlighted. The RubPro is divided into 3 sections, ‘fingers’ in lavender, ‘palm’ in marine blue and ‘thumb’ in teal. The catalytic residues are coloured in orange, Zn^2+^-coordinating residues in green and Zn^2+^ as red spheres. (**B**) Model of RubPro with a right-hand architecture, from the front-facing the catalytic site. A view of RubPro with a 120° lateral rotation is also shown, with the N- and C-terminals labelled. (**C**) Atomic view of Zn^2+^-coordination sites and the catalytic site. The distances are shown as black dashed lines in Å. (**D**) Two-dimensional topology map of the RubPro model, showing α-helices as cylinders and β-sheets as arrows. α-helices and β-sheets are labelled and numbered starting from the N-terminal. (**E**) Sequence alignment of RubPro with RUHV and RUSV proteases. A perfect alignment of residues is indicated in red with a (*), while alignment of residues with very strong similarities is indicated in orange with a (:). The catalytic dyad is boxed in blue, and the Zn^2+^-coordinating sites are boxed in green.

Our structure also confirmed Liu et al.’s findings that C1175, C1178, C1225 and C1227 are Zn^2+^-binding cysteines [30]. A previously unknown quartet of Zn^2+^-binding residues, C1034, C1037, C1040 and H1088, were also uncovered at the N-terminal of this novel structure. These two Zn^2+^-coordinating sites were well-defined by the characteristic tetrahedral coordination geometry of sulfur and nitrogen atoms from Cys and His respectively. At the first Zn^2+^ binding site, the residues C1034, C1037, C1040 and H1088 are located 2.4 Å, 2.3 Å, 2.4 Å and 2.1 Å away from Zn^2+^. While at the second Zn^2+^ binding site, all cysteine residues are 2.3 Å away from Zn^2+^ (Fig 2C).

The catalytic dyad C1152 and H1273 are located on the palm and the thumb respectively. The positions of these catalytic residues are similar to that of SARS-CoV-2 PLpro, whereby the catalytic triad is also found at the interface between the palm and the thumb regions [66]. The sulfur atom of C1152 is 4.2 Å apart from the delta nitrogen of H1273 (Fig 2C). In contrast to other cysteine proteases, no stabilizing Asn/Asp residue was found near H1273.

Sequence alignment of the RubPro with the protease domain of RUHV and RUSV was performed to analyse for the presence of conserved motifs among the *Rubivirus* genus. The protease domain of RUHV and RUSV shows 53 % and 40 % amino acid similarity respectively to RubPro, with conserved catalytic dyad and zinc coordinating residues (Fig 2E).

### Structural alignment with other cysteine proteases

The Dali server was utilised to search for structural homologs in the PDB database (Table S2) [67]. We selected the crystal structures of human ubiquitin specific protease (USP) 30 (PDB: 5OHN) [68], FMDV PCP (PDB: 4QBB) [69] and SARS-CoV-2 PLpro (PDB: 6WZU) [66] for further analysis against that of RubPro (Fig 3). Similar structural motifs might provide insights into protease functions and targets of the viral protease.

**Figure 3.**
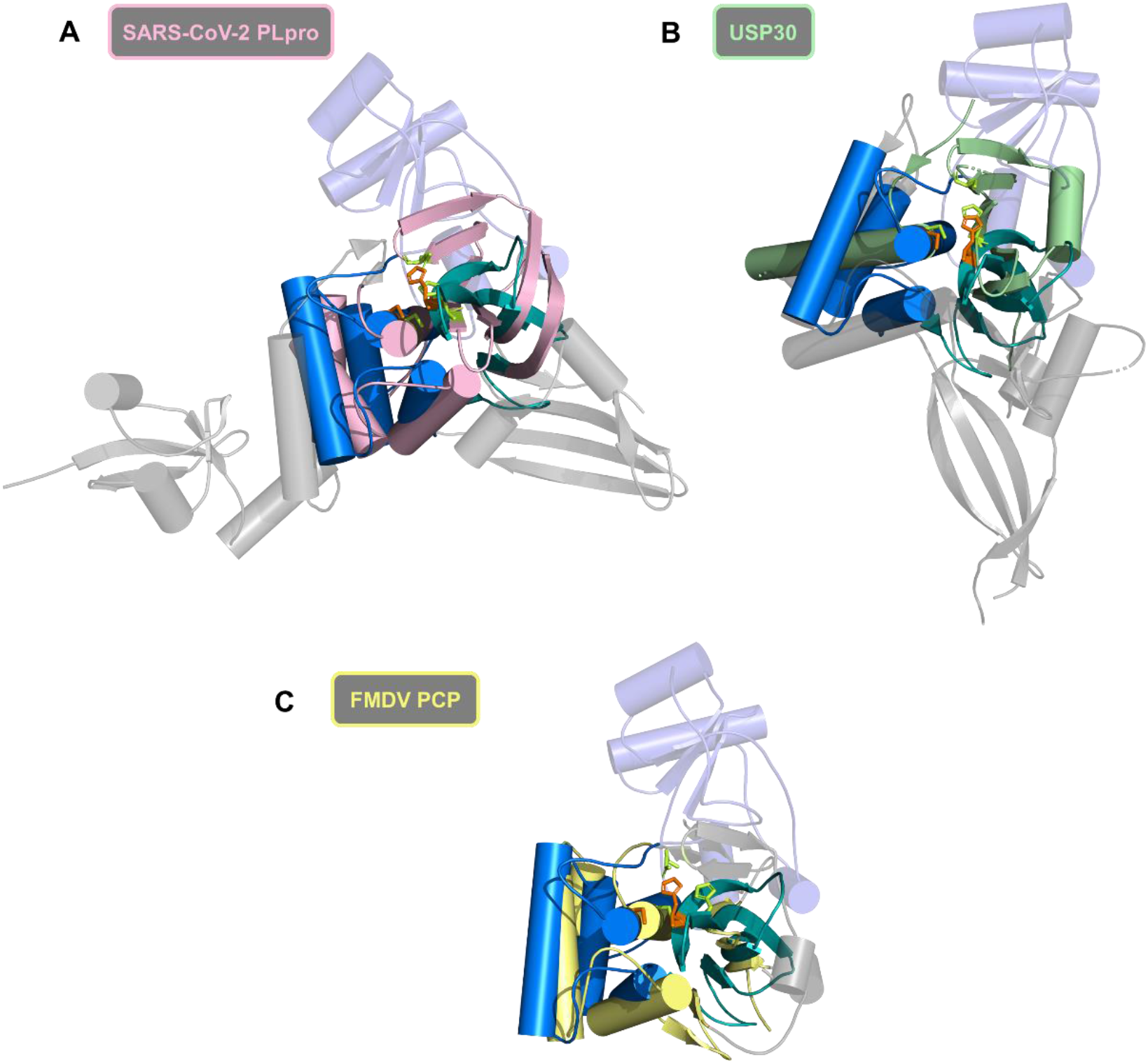
Structural alignment of RubPro with Human USP30, FMDV PCP and SARS-CoV-2 PLpro. RubPro follows the same colouring scheme as in Fig. 2, with its distinct N-terminal fingers domain depicted in transparent lavender cartoon. Distinct domains of the comparison proteases, that are not aligned with RubPro, are depicted in transparent grey cartoons. Catalytic residues of the comparison proteases are shown as lime sticks. (**A**) Residues 58-97, 107-175 and 422-495 of USP30 were aligned with RubPro, depicted in pale green. (**B**) Residues 43-126, 137-153 and 177-187 of FMDV PCP were aligned with RubPro, depicted in light yellow. (**C**) Residues 103-176 and 241-306 of PLpro were aligned with RubPro, depicted in pink.

For all 3 cysteine proteases, there was a good alignment of the core domains around the catalytic sites. The catalytic residues were in similar positions with less than 5 Å deviations (Fig S2). While RubPro has a Cys-His catalytic dyad, the other proteases had catalytic triads; Cys-His-Asp for PLpro and FMDV PCP, and Cys-His-Ser for USP30. The N-terminal fingers domain of RubPro was distinct and had no structural homologs with any of the other 3 proteases.

### RubPro cleaves RubProHel C1152A

RubProHel C1152A, with the natural cleavage junction between RUBV protease and helicase, was used to assess the trans cleavage activity of the RubPro. A catalytic C1152A mutation was introduced to RubProHel, to prevent any self-cleavage in cis. The RubProHel C1152A was incubated with doubling concentrations of RubPro. The in *vitro* trans cleavage activity of RubPro was demonstrated by the decreased of RubProHel C1152A, 64.9 kDa, and the formation of the helicase domain, 34.7 kDa (Fig 4A).

**Figure 4.**
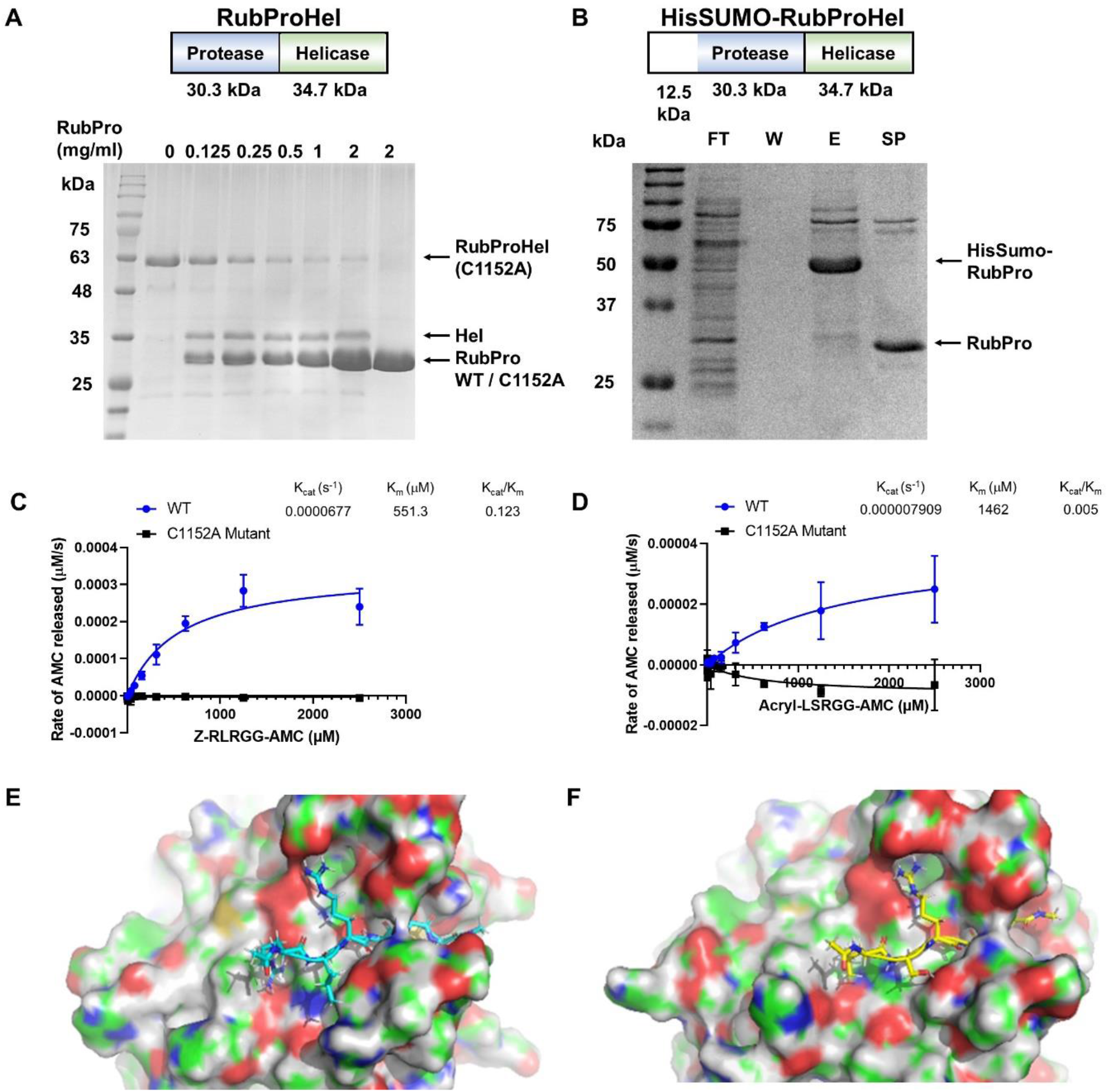
Enzymatic activity of RubPro. (**A**) SDS-PAGE profile of in vitro trans cleavage activity of RubPro using RubProHel C1152A as a substrate. 0.5 mg/mL of RubProHel C1152A was incubated with 0.125 mg/mL to 2 mg/mL of RubPro, at room temperature for 18 h in buffer (25 mM HEPES buffer at pH 7.5, 150 mM NaCl, 5% w/v glycerol and 2 mM DTT). The molecular weight of the RubPro is 30.3 kDa and RubProHel is 64.9 kDa. While the molecular weight of the expected cleavage products helicase and RubPro are 34.7 kDa and 30.3 kDa respectively. (**B**) Expression of HisSUMO-RubProHel WT (77.5 kDa) in *E. coli*. It is cleaved into HisSumo-RubPro (42.8 kDa, in the elution E) and helicase (34.7 kDa), which is not apparent in the flowthrough. RubPro (30.3 kDa) is obtained after Sumo protease cleavage (lane SP), validating the identity of the 42.8 kDa protein. The protease activity of RubPro was measured using the (**C**) Z-RLRGG-AMC and (**D**) Acryl-LSRGG-AMC substrates. Assays were carried out as triplicates at 37 °C at 5 μM enzyme concentration with varying substrate concentrations ranging from 0 to 2.5 mM. (**E**) The complex of the protein and the peptide RLRGGG. (**F**) The complex of the protein and the peptide LSRGGG.

### RubProHel is cleaved during expression

Expression of RubProHel in *E. coli* cells was used to ascertain cleavage of the p150-p90 junction. RubProHel is a construct with RUBV helicase at the C-terminal of RubPro. RubProHel WT and a C1152A mutant, both with an N-terminal His-Sumo tag, were expressed and purified. From Fig 4B, the WT expression of His-Sumo-RubProHel produces the cleaved products, the N-terminal 42.8 kDa His-Sumo-RubPro and the 34.2 kDa helicase at the C-terminal. The identity of the N-terminal cleavage product was validated by Sumo protease cleavage, which cleaved the 42.8 kDa His-Sumo-RubPro into 30.3 kDa RubPro. This shows that there is protease activity in RubProHel, whether in *cis* or *trans*, in *E. coli* expression. A mutant of the catalytic dyad, C1152A, showed no cleavage of RubProHel (Fig S3).

### Enzymatic activity of RubPro on two peptide substrates

Enzyme concentration and buffer pH optimisation was first conducted to identify the suitable conditions for conducting protease assay. Enzyme concentration greater than 3.75 μM was chosen for easier quantification and calculation of the enzymatic rate (Fig S4A). Buffer pH 6.4 to 6.6 displayed the highest enzymatic rate (Fig S4B). Finally, the protease activity of RubPro was measured at a protein concentration of 5 μM at 50 mM potassium phosphate buffer pH 6.6.

The protease assay of RubPro and RubPro C1152A mutant were performed using Z-RLRGG-AMC substrate and Acryl-LSRGG-AMC substrate. The RubPro has a higher affinity towards Z-RLRGG-AMC substrate than Acryl-LSRGG-AMC substrate with K_m_ values of 580 μM and 1460 μM respectively (Fig 4C and Fig 4D). The RubPro also exhibits a higher catalytic rate of 0.0000741 s^-1^ against Z-RLRGG-AMC substrate while having a 10-fold slower catalytic rate of 0.00000791 s^-1^ against Acryl-LSRGG-AMC substrate (Fig 4C and Fig 4D).

### Binding of the two substrate peptides to the RubPro active site

Two blocked hexamer peptides, Ace-Leu-Ser-Arg-Gly-Gly-Gly-Nme (LSRGGG) and Ace-Arg-Leu-Arg-Gly-Gly-Gly-Nme(RLRGGG), were simulated together with the Rubella virus protease. Several steps which combined peptide docking, molecular dynamics (MD) simulations and manually structure manipulations, were needed to find the optimized poses of the peptides. The details are provided in the Method part. During the final 100 ns MD simulations, the root mean squared deviation (RMSD) of the peptides after superimposing the protease showed stable binding (Fig S5). The RMSD values of LSRGGG have larger fluctuations than those of RLRGGG, indicating a bit less stable binding of LSRGGG than RLRGGG. The binding modes of the two peptides are quite similar: the last three glycine residues sit in the narrow groove formed mainly by residues: G1213, W1153 and C1152C on one side and T1271, G1272 and H1273 on the other side. The common third residue, Arginine, forms salt bridges with D1206 and D1217, showing the arginine residue is preferred here to anchor the peptide around the active side.

The side chain of serine residue of LSRGGG forms hydrogen-bond with the side chain of T1271. The first residue leucine has hydrophobic contacts with L1216, L1247 and L1291 which form a hydrophobic patch. However, such interaction is not very stable, during the simulation, the side chain of this leucine was found to flip out and the methyl group of Ace makes close contacts with this hydrophobic patch.

The first residue, arginine, of RLRGGG, seems more fit in this position. Its side chain not only makes contacts with the hydrophobic patch (L1216, L1247 and L1291) on one side; on the other side stacks with the ring of H1290, also makes hydrogen bond with the carbonyl group of H1290 and a salt-bridge with D1214. The second residue, leucine, however, seems no good interaction partners, loosely interacts with the methyl group of T1271.

The MMGBSA method was used to calculate the binding energy between the protein and the peptides. The results are consistent with the simulation RMSD values: the peptide RLRGGG has a binding energy of -49.06± 4.68 kcal/mol and the peptide LSRGGG has -42.76±7.13 kcal/mol. The slightly stronger binding of RLRGGG is also in line with the experimental measured K_m_ values.

### RubPro is a K48-linkage-specific deubiquitinase

To determine Rubpro activity on Ubiquitin (Ub) and the Ub-like (Ubl) modifier interferon stimulated gene 15 (ISG15), we purified K63-triubiquitin, K48-triubiquitin and M1-triubiquitin(linear) chains and ISG15-HIS according to our previous work [65]. Starting with ubiquitin, robust RubPro activity and high specificity was observed towards Lys48-linked polyubiquitin (Fig 5B), where triubiquitin was cleaved to monoubiquitin products, but not Lys63-linked triubiquitin (Fig 5A) and linear-linked triubiquitin (Fig 5D). Similar results were obtained using gel-based analysis of Lys63 and linear-triubiquitin versus cleavage of ISG15-HIS to mature ISG15, HIS tag is not removed from the ISG15 C-terminus (Fig 5C). A mutant of the catalytic dyad, C1152A, showed no cleavage of K63-triubiquitin, K48-triubiquitin and M1-triubiquitin (linear) chains and ISG15-HIS (Fig 5A-D). Time course analysis also suggest that RubPro is a K48-linkage-specific deubiquitinase (Fig 5E).

**Figure 5.**
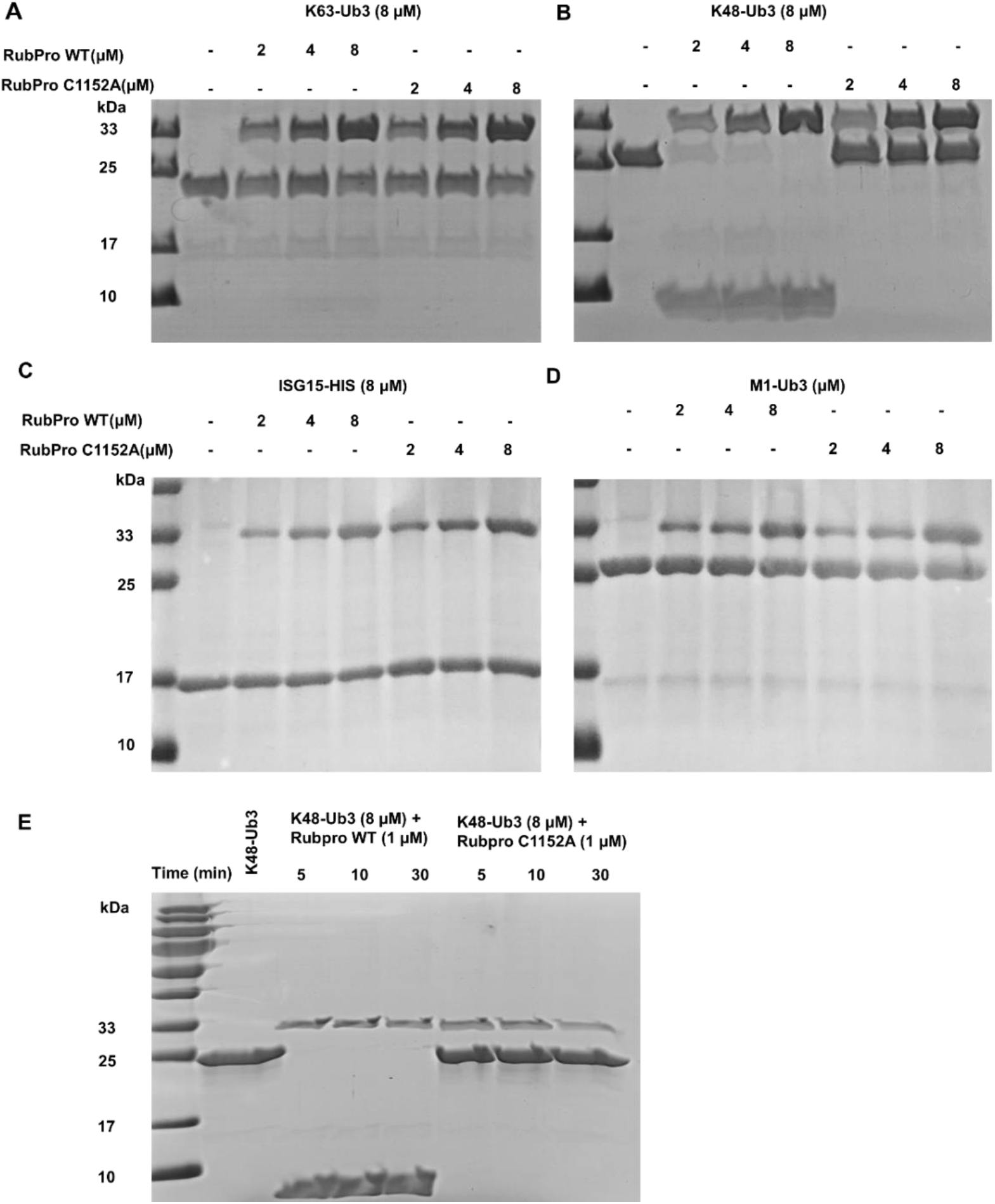
RubPro is a K48-Ubi specific deubiquitinase. Wild-type and C1152A mutants of RubPro were tested for their ability to process triubiquitin and ISG15 in vitro. Purified RubPro at indicated concentration were incubated at 25°C with K63-Ub3(**A**),K48-Ub3(**B**),ISG15-HIS(**C**), M1-Ub3(**D**) in a reaction buffer for 10min. Cleavage products were analyzed by SDS-PAGE and stained with Coomassie blue. (**E**) Time course analysis of triubiquitin (8 μM) hydrolysis using 1 μM RubPro in a reaction buffer, resolved on a Coomassie-stained SDS–PAGE gel.

## Discussion

The novel structure of RubPro presented here sheds much insight into the poorly understood RUBV (Fig 2). The novelty of the structure is further exemplified by the unsuccessful prediction using the Alphafold2 server (Fig S6). This 1.64 Å structure represents the first structure of the RUBV non-structural protein. All 10 of the other PDB entries of RUBV proteins are the structural E1 glycoprotein and capsid proteins. While RubPro is similar to other PCPs, especially at the domains surrounding the catalytic dyad, it has a very distinct and unique N-terminal ‘fingers’ domain. The domain can be coordinating the Zn^2+^ cation for structural support. It may also serve as a docking site for other cleavage targets. More work needs to be done to better understand and characterise this unique N-terminal domain.

In contrast to the previous study [34], we did not identify the presence of Ca^2+^-binding sites nor a EF-hand domain in our crystal structure. We tried supplementing CaCl_2_ into the crystallisation buffer and soaking the crystals in a solution containing CaCl_2_. One possible future work would be to conduct a denaturation-refolding assay of RubPro in the presence of Ca^2+^. One surprise of the RubPro structure is that the C-terminal ends in a β-sheet β8 that is pointing away from the catalytic site (Fig 2B). This suggests that the cleavage junction SRGG/GTCA in the p200 polypeptide could be inaccessible to the RubPro catalytic domain, for *cis* cleavage of the p150-p90 junction. A possibility could also be RubPro might function like the Chikungunya virus capsid protease domain, which is only active for one proteolytic reaction, after which the active site is inaccessible [70]. Structural work needs to be carried out to solve the structure of RubProHel, to shed light on this apparent structural disagreement. Co-crystallisation can also be carried out with a bound inhibitor or a substrate to further characterise the active site. A C1152A or H1273A mutant could be used to permanently bind a substrate without cleaving it.

The newly discovered *Rubivirus* members, RUHV and RUSV, are the closest relatives of RUBV. From sequence alignment of RubPro with RUHV and RUSV proteases (Fig 2E), there are conserved motifs, especially around the catalytic dyads of C1152 and H1273. Unlike RUBV, RUHV and RUSV are pathogens in non-human mammals, capable of crossing host species barriers. It is not shown that humans can be infected with RUHV or RUSV, but there are some concerns about future zoonoses arising from RUBV-like viruses like RUHV and RUSV. Knowledge about RubPro may prove useful for other *Rubivirus* proteases given that conservation of the important motifs is observed.

Despite limited sequence similarity with other cysteine proteases, RubPro is structurally similar to the other cysteine proteases, namely, human USP30, FMDV PCP and SARS-CoV-2 PLpro at the catalytic core (Fig 3 and Table S2). The structure of RubPro was also compared with the protease domain of the Chikungunya virus, as RubV was previously classified under the Togaviridae family. However, both structures are distinctive different (Fig S7), and no members of the Togaviridae family were identified as hits in the Dali structural homology database search (Table S2).

Expression of the WT RubProHel showed significant cleavage into the 42.8 kDa and 34.7 kDa products (Fig 4B), demonstrating *in vivo* protease activity in *E. coli* cells. As such, the RubProHel C1152A mutant is likely to be a good candidate for structural studies, as it can bring structural insights into the self-cleavage mechanism of p200 into p150 and p90. The orientation of the cleavage junction SRGG/GTCA relative to the catalytic site can inform on the mechanism of RubPro activity on p200, whether it works in *cis* or *trans* dominantly. RubProHel C1152A was cleaved by RubPro in trans (Fig 4A), and this differs from Liang et al.’s conclusion that *trans* cleavage requires additional residues 920-974 [20]. The additional residues might contribute to improved protease activity but is not essential for protease activity. In our study, RubPro at a high concentration of 2 mg/mL or 66 μM, was unable to fully process all the RubProHel C1152A substrate, which is at a lower concentration of 0.5 mg/mL or 7.7 μM, at room temperature after 18 h. Furthermore, the RubPro also displayed weak enzymatic activity having K_cat_ of 0.000074 and affinity for the Z-RLRGG-MCA substrate with K_m_ of 580 μM. A RubPro construct containing the upstream residues from 920 onwards, could perhaps display more active trans cleavage activity.

Given the structural similarities to numerous USPs, FMDV protease and SARS-CoV-2 PLpro (Fig 3 and Table S2), we seek to investigate if RubPro also demonstrates deubiquitination activity. RubPro was shown to be a K48-linkage-specific deubiquitinase (Fig 5). Similar K48-linkage deubiquitination was observed for Herpes Simplex Virus 1 ubiquitin-specific protease and Porcine Reproductive and Respiratory Syndrome Virus nonstructural protein 2, whereby the removal of K48-Ub from IκBα, prevents the degradation of IκBα and the activation of both NF-κB signalling pathway and type I interferon response [71, 72]. Future works seek to investigate the cellular targets of RubPro and the possible role of RubPro in modulating innate immune responses during RUBV infection.

## Conclusion

Given the lack of fundamental understanding of RUBV replication processes, we set out to characterise the functional protease domain of the non-structure p200 and p150 polypeptides, both structurally and functionally. RubPro might be a drug target candidate as an antiviral therapy for RUBV infections. Here, we managed to express and purify a pure RubPro construct and capture high-quality crystals for x-ray diffraction. This led to the structure of RubPro which we solved at 1.64 Å resolution. This novel structure provides much insight into RubPro, but by the same token, poses many new questions which are yet to be answered. We have validated the Zn^2+^-binding sites atomically and structurally. We found a strong structural alignment of RubPro with SARS-CoV-2 PLpro and FMDV PCP at the core regions around the catalytic sites. Sequence alignment of RubPro with RUHV and RUSV proteases show strong conservation of certain motifs. Overall, we have successfully characterised the structure of RubPro, but our construct has relatively low protease enzymatic activity. We were able to demonstrate cleavage of RubProHel in *E. coli*, and RubPro showed *trans* protease activity against the natural substrate RubProHel C1152A mutant. Host factors that are targeted by similar viral PCPs might be deubiquitinated by RubPro as well. The work presented here may prove to be merely the cusp of deeper research into RUBV replication.

## Acknowledgements

This research is supported by the Ministry of Education, Singapore, under its MOE AcRF Tier 1 Award 2021-T1-002-021. We wish to acknowledge the funding support for this project from Nanyang Technological University under the URECA Undergraduate Research Programme. J. P. Q. is supported by the Nanyang Presidential Graduate Scholarship and the Lee Kong Chian School of Medicine. We gratefully acknowledge the beamline staff at MXII beamline in Australian Light Source, Melbourne, Australia for providing us with outstanding support during the data collection.

