## Supplementary File for "Crystal structure of the Rubella virus protease reveals a unique papain-like protease fold"

### SUPPORTING INFORMATION

**Table S1**. X-ray data collection and refinement statistics (PDB code: 7FAV).

| **Data collection statistics** | |
|---|---|
| Wavelength (Å) | 1.284 |
| Resolution range (Å) | 41.37 - 1.64 (1.699 - 1.64) |
| Space group | P 43 21 2 |
| Unit cell a, b, c | 60.2, 60.2, 173.8 |
| a, b, g (Å) (°) | 90, 90, 90 |
| Total reflections | 2075865 (189870) |
| Unique reflections | 40150 (3811) |
| Multiplicity | 51.7 (49.8) |
| Completeness (%) | 99.67 (96.87) |
| Mean I/sigma (I) | 41.82 (4.42) |
| Wilson B-factor (Å$^2$) | 22.15 |
| [a]$R_{merge}$ | 0.08663 (1.09) |
| R-meas | 0.08749 (1.101) |
| R-pim | 0.01207 (0.1525) |
| CC1/2 | 1 (0.921) |
| CC* | 1 (0.979) |
| **Refinement statistics** | |
| Reflections used in refinement | 40150 (3811) |
| Reflections used for R-free | 2074 (185) |
| [b]$R_{work}$ | 0.1801 (0.2163) |
| [c]$R_{free}$ | 0.1988 (0.2083) |
| Number of non-hydrogen atoms | 2289 |
| macromolecules | 2073 |
| ligands | 44 |
| solvent | 172 |
| Protein residues | 271 |
| [d]RMSD (bonds) (Å) | 0.007 |
| RMSD (angles) (°) | 1.19 |
| Ramachandran favored (%) | 98.88 |
| Ramachandran allowed (%) | 1.12 |
| Ramachandran outliers (%) | 0.00 |
| Rotamer outliers (%) | 0.00 |
| Clashscore | 4.81 |
| Average B-factor | 27.77 |
| macromolecules | 26.32 |
| ligands | 43.47 |
| solvent | 41.13 |
| Number of TLS groups | 7 |

Statistics for the highest-resolution shell are shown in parentheses.
[a]$R_{merge} = \sum|I_j - < I >|/\sum I_j$, where $I_j$ is the intensity of an individual reflection, and $< I >$ is the average intensity of that reflection.
[b]$R_{work} = \sum||F_{obs}| - |F_{calc}||/\sum|F_{obs}|$, where $F_{obs}$ denotes the observed structure factor amplitude, and $F_c$ is the structure factor amplitude calculated from the model.
[c]$R_{free}$ is as for $R_{work}$ but calculated with 5% of randomly chosen reflections omitted from the refinement.
[d]RMSD, root mean square deviations.

**Table S2**. Selected top hits from Dali server output

| PDB | Protein description | Dali Z-score[a] | Sequence identity % | RMSD[b] / Å |
|---|---|---|---|---|
| 5OHN | Human ubiquitin-specific protease 30 (USP30) | 8.1 | 13 | 3.6 |
| 4QBB | Foot-and-mouth-disease virus leader protease (FMDV PCP) | 7.7 | 10 | 3.3 |
| 5K1B | Human ubiquitin-specific protease 12 (USP12) | 7.4 | 13 | 3.0 |
| 5L8H | Human ubiquitin-specific protease 46 (USP46) | 7.1 | 14 | 3.4 |
| 5TXK | Human ubiquitin-specific protease 35 (USP35) | 7.1 | 10 | 3.3 |
| 6WZU | SARS-CoV-2 papain-like protease (SARS-CoV-2 PLpro) | 5.4 | 11 | 3.7 |

[a]Z-score is an indicator of significant similarity.

[b]RMSD, root mean square deviations

**Figure S1**. An anomalous difference Fourier map is shown as black mesh at 6.0 σ. RubPro follows the same colouring scheme as in Fig. 2. The 'fingers' coloured in lavender, 'palm' in marine blue and 'thumb' in teal. $Zn^{2+}$ are represented as red spheres.

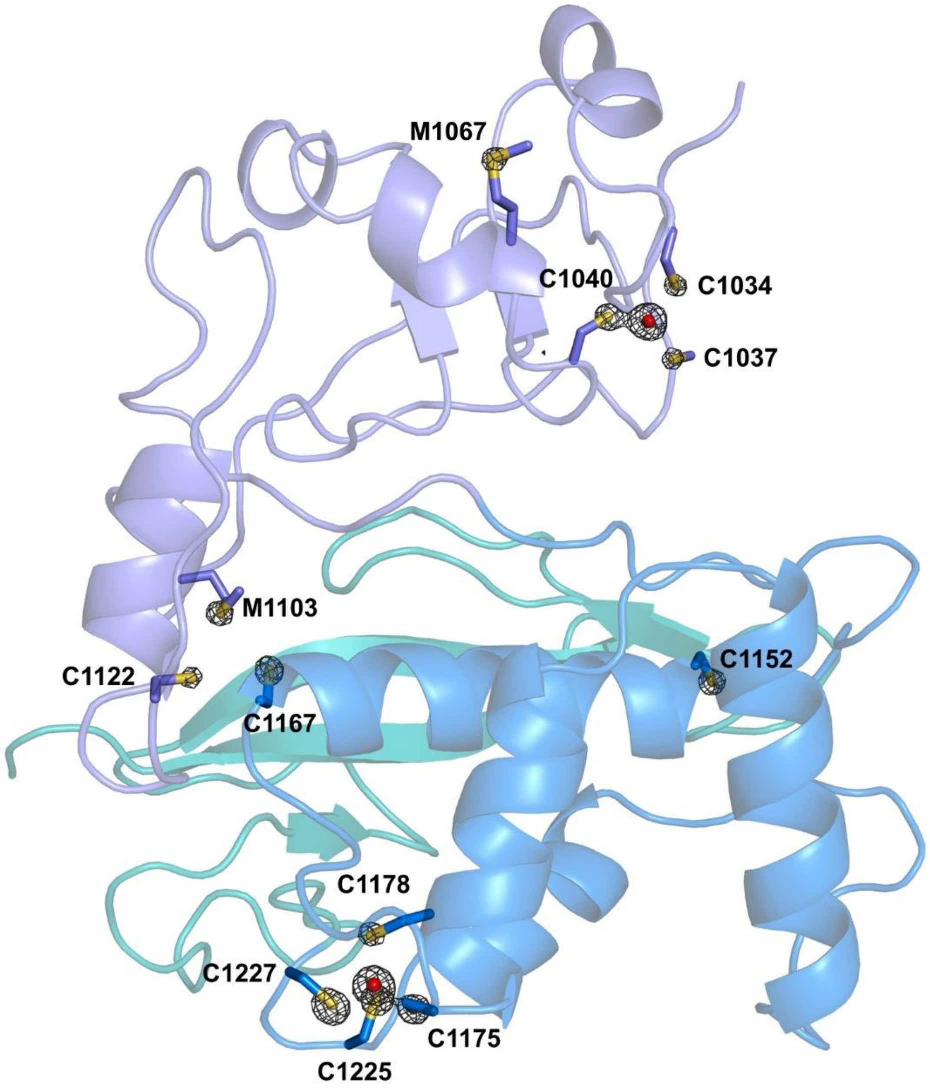

**Figure S2**. Closed up view of the catalytic residues of Rubpro with the superimposed human USP30 (Left panel), FMDV PCP (Middle panel) and SARS-CoV-2 PLpro (Right panel). The distances are shown as black dashed lines in Å.

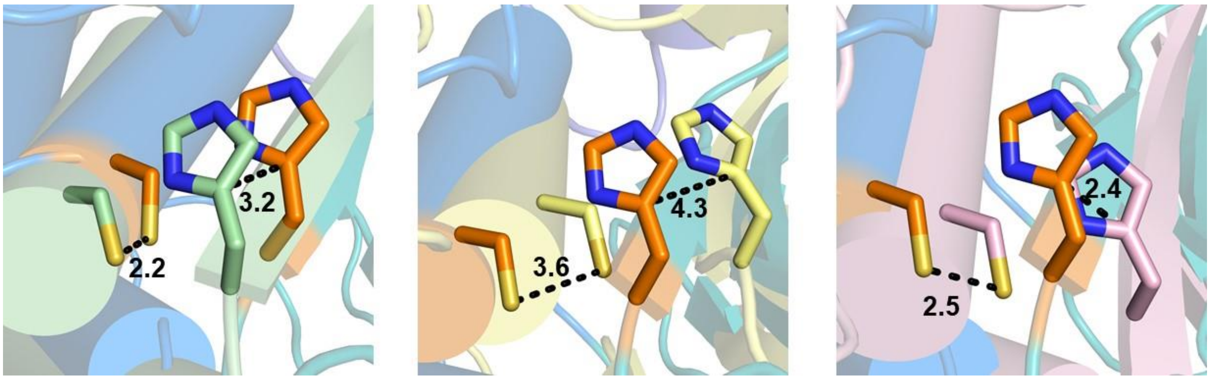

**Figure S3**. (A) Expression of RubProHel C1152A (77.5 kDa) in E. coli. It remains uncleaved as His-Sumo-RubProHel (77.5 kDa) (lanes E1 and E2). (B) RubProHel (64.9 kDa) is obtained after Sumo protease cleavage (lane SP), validating the identity of the 77.5 kDa protein

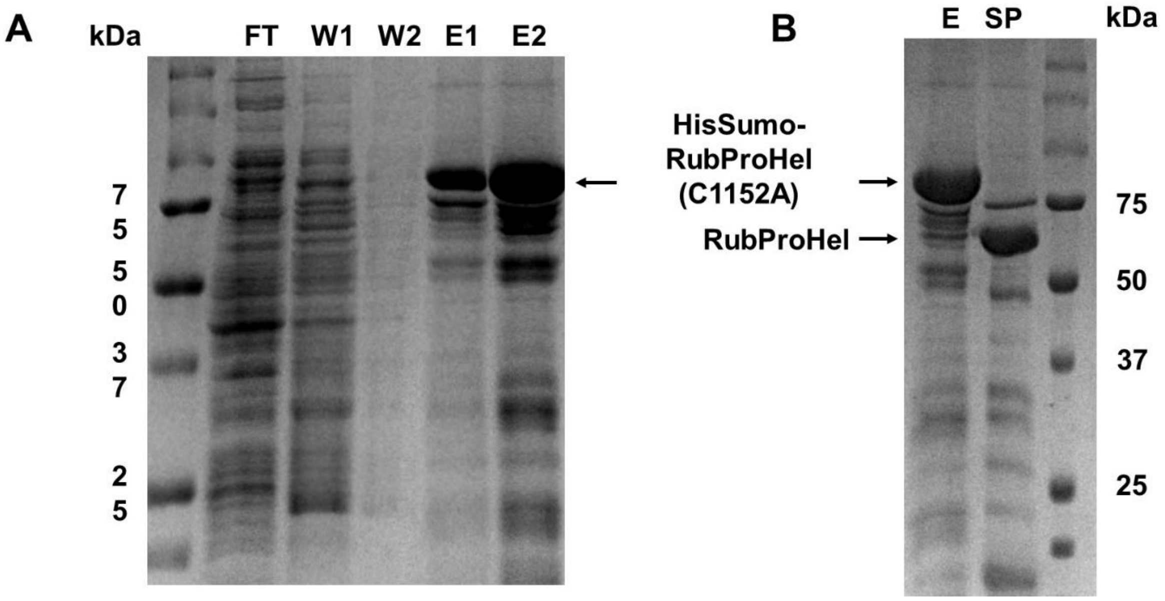

**Figure S4**. (**A**) Enzyme concentration optimisation of RubPro. The RubPro were screened from enzyme concentration of 0.93 μM to 15 μM using 50 μM of Z-RLRGG-AMC. (**B**) Buffer pH optimisation for protease assay using 5 μM of RubPro and 50 μM of Z-RLRGG-AMC from pH 5.8 to 7.8.

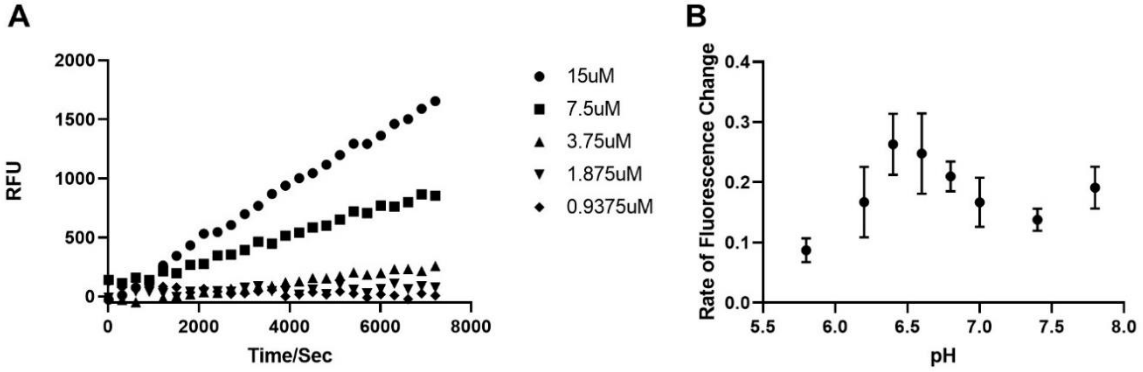

**Figure S5**. The root mean squared deviation (RMSD) of the peptides after superimposing the protein.

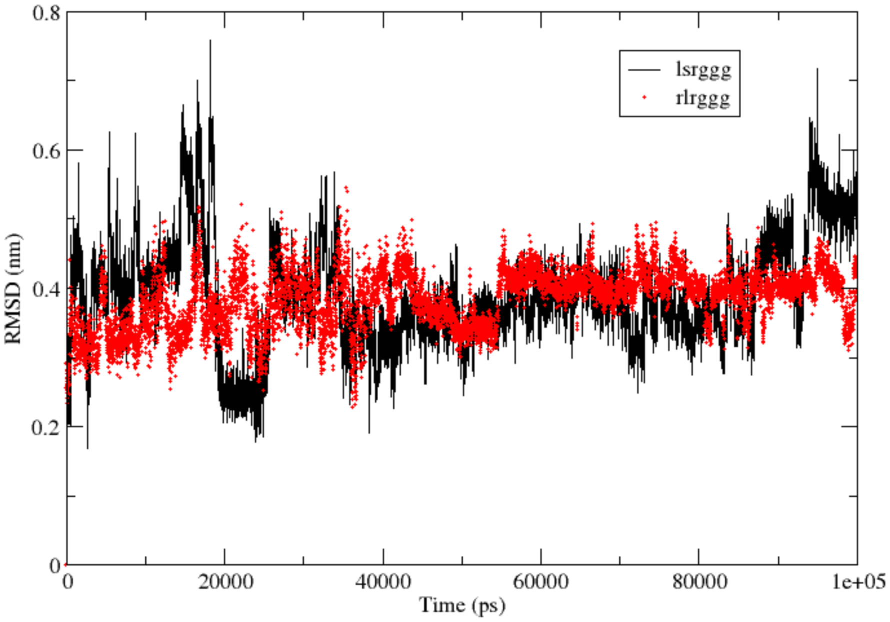

**Figure S6**. Comparison of the crystal structure of RubPro with Alphafold2's structural prediction. The solved crystal structures of RubPro (PDB: 7FAV) is coloured in red. The relaxed models generated from Alphafold2 are coloured in green, magenta, yellow, pink and grey.

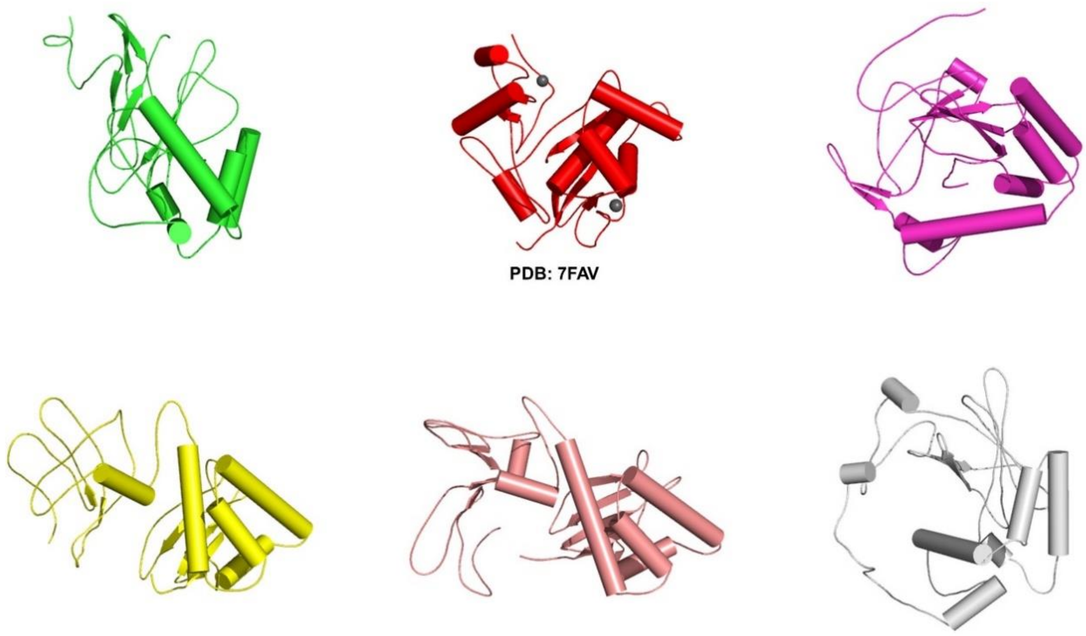

**Figure S7**. Comparison of the crystal structure of RubPro with protease from alphavirus genus. The RubPro is shown on the left panel and the Chikungunya protease (PDB ID: 4ZTB) is shown on the right panel. The location of the catalytic site is highlighted in magenta.

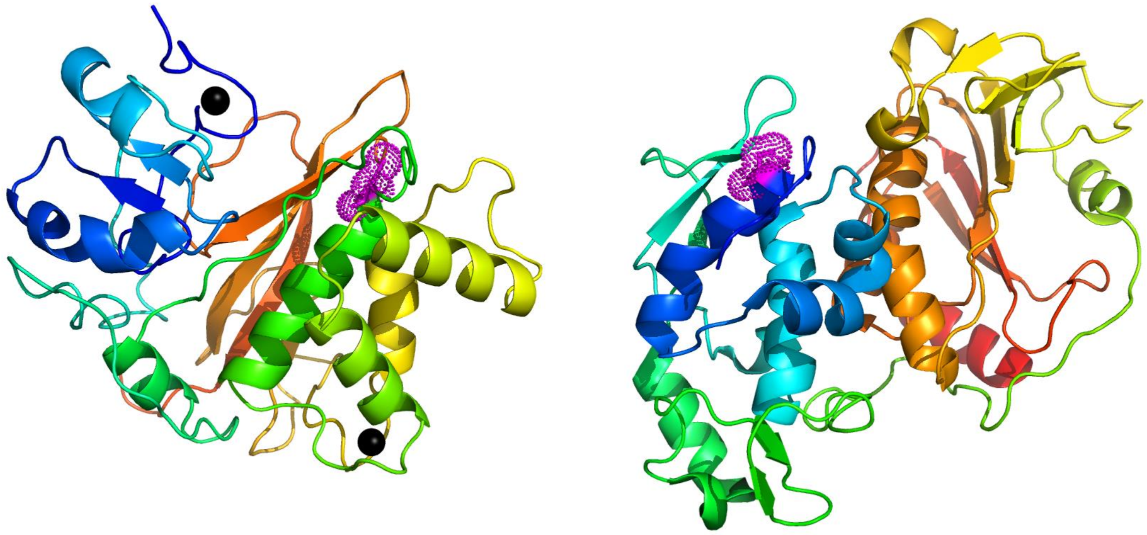
